# Sex-Specific Complement and Cytokine Imbalances in Drug-Resistant Epilepsy: Biomarkers of Immune Vulnerability

**DOI:** 10.1101/2024.09.16.612934

**Authors:** Nicole Pinzon-Hoyos, Yibo Li, Monnie McGee, Nicholas P. Poolos, Nicola Marchi, Amy L. Brewster

## Abstract

**Objective:** Drug-resistant epilepsy (DRE) poses significant challenges in treatment and management. While seizure-related alterations in peripheral immune players are increasingly recognized, the involvement of the complement system, central to immune function, remains insufficiently explored in DRE. This study aimed to investigate the levels of complement system components and their association with cytokine profiles in patients with DRE.

**Methods:** We analyzed serum samples from DRE patients (n = 46) and age- and sex-matched healthy controls (n = 45). Complement components and cytokines were quantified using Multi- and Single-plex ELISA. Statistical analyses examined relationships between complement molecules, cytokines, and clinical outcomes including epilepsy duration, Full-Scale Intelligence Quotient (FSIQ) scores, and age.

**Results:** We found common alterations in all DRE cases, including significant complement deficiencies (C1q, Factor H, C4, C4b, C3, and C3b/iC3b) and detectable bFGF levels. DRE females showed significantly lower levels of TNFα and IL-8 compared to healthy females. We observed a trend towards elevated CCL2 and CCL5 levels in DRE males compared to healthy males. These findings suggest potential sex dimorphism in immune profiles. Our analysis also indicated associations between specific complement and inflammatory markers (C2, IL-8, and IL-9) and Full-Scale Intelligence Quotient (FSIQ) scores in DRE patients.

**Interpretation:** Our study reveals sex-specific peripheral complement deficiencies and cytokine dysregulation in DRE patients, indicating an underlying immune system vulnerability. These findings provide new insights into DRE mechanisms, potentially guiding future research on complement and cytokine signaling toward personalized treatments for DRE patients.

## Introduction

Epilepsy is a chronic neurological disorder characterized by spontaneous and recurrent seizures. One-third of epilepsy patients experience seizures that are resistant to standard anti-seizure medications (ASM), a condition known as drug-resistant epilepsy (DRE) ^1^. Within the multifaceted causation of DRE, contemporary studies indicate the presence of immune system dysregulations with a complex interplay between brain and peripheral inflammation, destabilizing the neurovascular interface and possibly contributing to ictogenesis ^2-5^. Cellular and molecular immunity players have been examined in blood in association with seizures and epileptic conditions ^3, 6-9^, with indications of altered levels of pro-inflammatory cytokines such as Interleukin 6 (IL-6), IL-1β, and Tumor necrosis factor alpha (TNFα) suggesting increased inflammation ^10-15^.

This emerging research has, however, neglected the immune complement system, a key orchestrator of inflammation ^16^. The complement system consists of over 30 small proteins produced in the liver and found in blood and tissue fluids ^16-18^. Under healthy conditions, the complement system contributes to immune surveillance and homeostasis. Its activation occurs through three pathways: classical, lectin, and alternative (Fig. 1A), triggering proteolytic reactions that generate complement proteins. These proteins enhance the immune response by acting as anaphylatoxins, chemokines, opsonins, and cell-lysis complexes ^16-18^. Activation of the complement system enables the release of inflammatory cytokines, which can, in turn, further activate the complement cascade, creating a feedback loop crucial for modulating immune responses ^16^. Complement system dysregulations occur in infections, autoimmune disorders ^19-21^, and neurological conditions ^22^, including epilepsy ^23-25^, potentially impacting symptoms and disease progression.

Given the complement system’s crucial role in immune regulation, investigating its activation in epilepsy is of significant interest, with possible diagnostic and therapeutic value. Studies on circulating complement levels in epilepsy patients have yielded mixed results ^23 24^. One report found lower levels of complement component iC3b and higher levels of C4 in patients with uncontrolled focal seizures ^23^, while another showed reduced C3 and C4 levels in patients with idiopathic generalized epilepsy ^24^. Gene ontology enrichment and KEGG pathway analyses of serum from patients with DRE revealed upregulation in complement activation and coagulation cascades ^25^. However, the extent of complement alterations and their relationship to the inflammatory response remains unclear. To address this knowledge gap, we measured the levels of multiple complement components and cytokines in the serum of individuals with DRE. We compared them to healthy age- and sex-matched controls. In this cohort, we assessed potential sex dimorphism in complement and cytokine serum concentrations and their relation to epilepsy duration, Full-Scale Intelligence Quotient (FSIQ) scores, and age.

## Methods

### Ethics Statement

Serum samples were collected with patients’ informed consent under approved Institutional Review Board (IRB) protocol #1011004282 (Development of a Biorepository for Methodist Research Institute; Indiana University Health Biorepository). All identifiable information was removed before conducting experiments and analyses under IRB protocol #21-126 (Southern Methodist University). Following collection, serum samples were stored at -80°C.

### Sample population

The study included serum from 46 patients with DRE (Females, n = 22; Males, n = 24) and 45 age and sex-matched healthy individuals (controls) without a history of immune/inflammatory conditions (Females, n = 21; Males, n = 24) (Table 1). Blood samples were collected from this cohort of DRE patients as part of routine preoperative procedures. These patients subsequently underwent surgical resection to remove epileptogenic foci. Patients with DRE underwent neuropsychological assessment with a qualified neuropsychologist administering the Wechsler Abbreviated Scale of Intelligence (WASI) test. FSIQ scores were estimated from the WASI test.

**Table 1.** Demographic and neurophysiological information of patients with drug-resistant epilepsy (DRE) and healthy individuals.

| Characteristics | DRE Females | DRE Males | Healthy Females | Healthy Males |
| --- | --- | --- | --- | --- |
| Number of patients | 22 | 24 | 21 | 24 |
| Age range (years) | 21-69 | 20-68 | 22-65 | 19-69 |
| Mean age (years) | 38.54±12.70 | 38.25±13.78 | 36.86±11.77 | 37.79±14.15 |
| Duration of epilepsy (years) | 20.85±12.54 | 18.56±13.86 | N/A | N/A |
| Mean FSIQ score | 86.92±11.69 | 89.50±13.21 | N/A | N/A |
| Patients with FSIQ scores | 12 | 16 | N/A | N/A |

### Enzyme-linked Immunoassay (ELISA)

Complement and cytokine levels were measured using various Milliplex and ELISA kits: Complement Panel 1 (HCMPEX1-19K): C2, C4b, C5, C5a, MBL; Complement Panel 2 (HCMP2MAG-19K): C1q, C3, C3b/iC3b, C4, Factor B, Factor H; Human C3a ELISA Kit (BMS2089); C5b-9 Singleplex Panel (EZS1M-140K); Bio-Rad Pro Human Cytokine Grp I panel 27-plex (M500KCSF0Y). Kits were processed according to manufacturers’ instructions with serum dilutions ranging from 1:100 to 1:20000. Samples below the detection limit were excluded from analysis, including 12 cytokines from the 27-plex kit.

### Statistical analysis

Complement and cytokine comparisons between two groups (DRE and healthy controls) were performed using unpaired t-tests with Welch’s correction (Figure 1 and Supplementary Figure 5, Supplementary Table 1). We assessed variance equality between the two groups using Levene’s test. As the test showed significant variance differences (p < 0.05), we applied Welch’s t-test to compare group means, with α set at 0.05 for all tests. For analysis of complement and cytokine blood levels across sex and health status (condition), a two-way analysis of variance (ANOVA) was used to assess the main effects of condition (healthy vs. DRE) and sex (male vs. female), as well as their interaction (Figures 2, 4; Supplementary Figure 6; Supplementary Table 2). Šídák’s multiple comparisons test was then applied for post-hoc pairwise comparisons between groups. These analyses were conducted using GraphPad Prism (version 10.1.2, GraphPad Software). Correlations were calculated using simple linear regression and Pearson correlation coefficient (r) analysis using R software (Version 4.4.0). The r value measures the strength and direction of the linear relationship between two variables, with values ranging from -1 to 1 (Figures 3 and 5). Correlation strength was interpreted as follows: Positive correlation: High (0.5 to 1.0), moderate (0.30 to 0.49), and small (0.10 to 0.29); Negative correlation: High (−1.0 to -0.5), moderate (−0.49 to -0.30), and small (−0.29 to -0.10). No correlation was defined as r = 0.

## Results

### Abnormal complement cascade components in DRE

The complement system consists of the classical (C1q), lectin (mannose-binding lectin; MBL), and alternative pathways which facilitate the cleavage of C3 into C3a and C3b (Fig. 1A). Subsequently, C3b facilitates C5 cleavage into C5a and C5b, with C5b initiating the formation of the membrane attack complex (MAC) (terminal pathway). C3a and C5a act as inflammatory signals, while C3b and C5b can also aid in phagocytosis. To assess the status of the complement system in DRE patients, we quantified the levels of these key components and compared them to those in healthy controls (Fig. 1; see Table 1 for details on patient cohorts). The overall profile of these complement cascades indicated a general decrease in the serum levels of several complement components in DRE patients compared to healthy individuals (Fig. 1B). Specifically, the components C1q, C4, C4b, C3, C3b/iC3b, C5, and regulatory Factor B and Factor H were significantly less abundant, by ∼30%, in DRE patients compared to healthy controls (p < 0.05, Fig. 1C-O). In contrast, the levels of MBL, C2, C3a, C5a, and C5b-9 did not differ significantly between the two groups. These findings suggest a deficiency in complement levels in the serum of DRE patients compared to age-matched healthy individuals.

**Figure 1.**
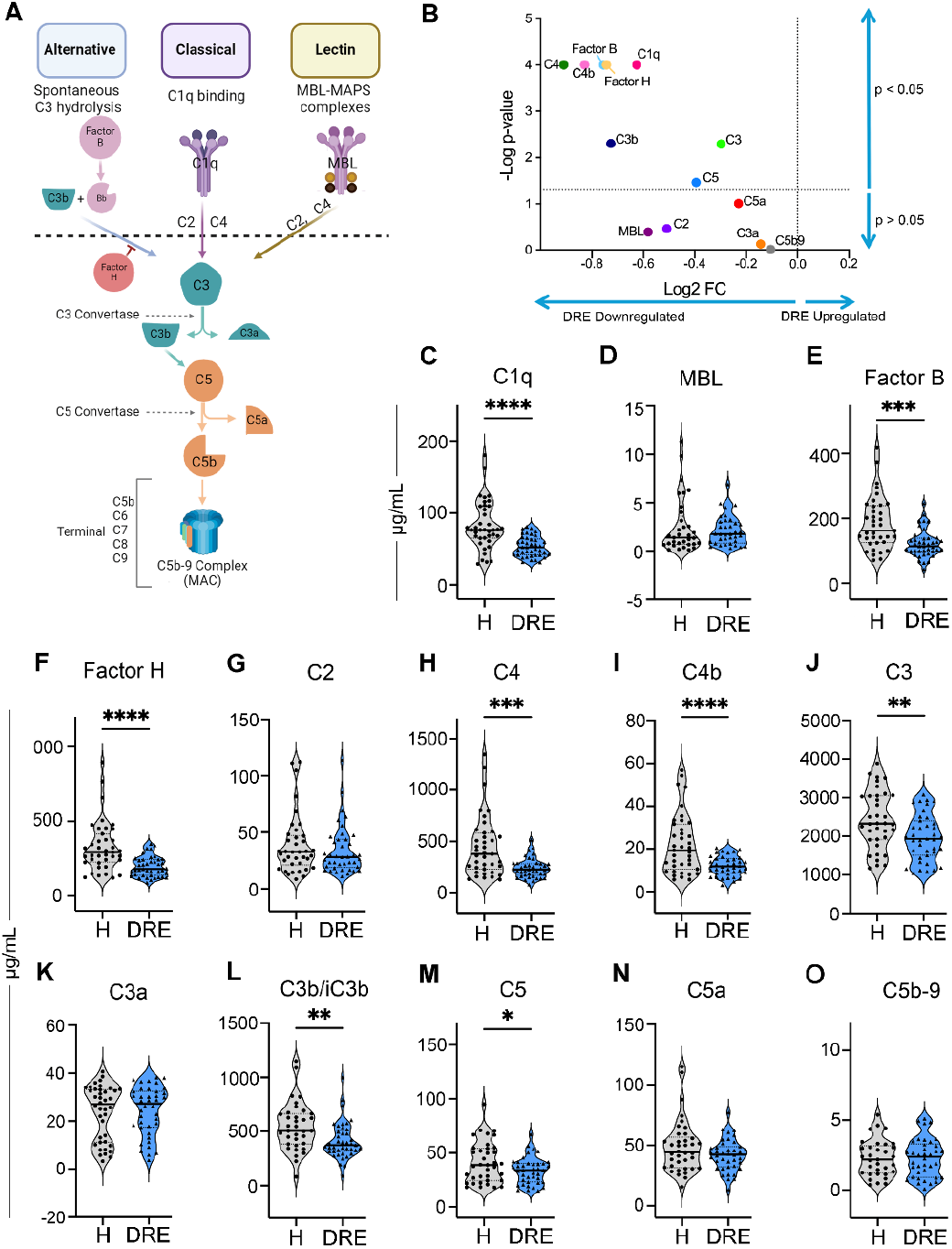
Serum complement component concentrations in patients with drug-resistant epilepsy (DRE) and healthy controls. **A**, The diagram illustrates the convergence of the three complement pathways: alternative, classical, and lectin, leading to the cleavage of C3 and C5. The alternative pathway is activated through spontaneous hydrolysis of C3. The formation of C3b and Bb components (from Factor B) allows the cleavage of C3, being regulated by Factor H. All the pathways converge in the cleavage of C5 into C5b which helps assemble the membrane-attack complex (MAC)(terminal pathway). **B**, A volcano plot showing the significance (unpaired *t*-test) and mean fold change of complement proteins between DRE patients and healthy controls. The X-axis represents Log2 Fold Change (FC), with dots on the left (< 0) indicating downregulated proteins and dots on the right (> 0) representing upregulated molecules. The Y-axis shows -Log p-value, with significant points appearing in the upper part of the plot; the higher the dots, the stronger the statistical significance. The significance threshold was set at 1.30103 (-log 0.05). **C-O**, Serum concentrations of multiple complement molecules in DRE patients (blue) and healthy controls (gray): C1q, Mannose-binding Lectin (MBL), Factor B, Factor H, C2, C4, C4b, C3, C3a, C3b/iC3b, C5, C5a, and C5b-9. Violin plots show individual data points, median, and quartiles. Statistical analysis was performed using an unpaired *t*-test. *p < 0.05, **p < 0.01, ***p < 0.001, ****p < 0.0001.

Next, we investigated whether sex-dependent differences existed (Fig. 2), as emerging evidence suggests that immune responses can differ between sexes ^26-32^. The levels of complement components were similar between healthy males and females (p > 0.05, Fig. 2A). (Two-Way ANOVA shown in Supplementary Table 2). A decrease of approximately 30% in the levels of complement components C1q, Factor B, C4, C4b, and Factor H was consistently observed in both female and male DRE groups compared to their healthy counterparts (p < 0.05, Figs. 2B and 2C). However, DRE males had significantly lower levels of C3b/iC3b compared to healthy males (C3b/iC3b, p = 0.039), while DRE females showed slight non-significant reductions (C3b/iC3b, p = 0.372). The C3a, C5, C5a, and C5b-9 levels were similar between males and females in both groups (p > 0.05). These findings indicate that males and females with DRE have deficiencies in the levels of circulating complement components.

**Figure 2.**
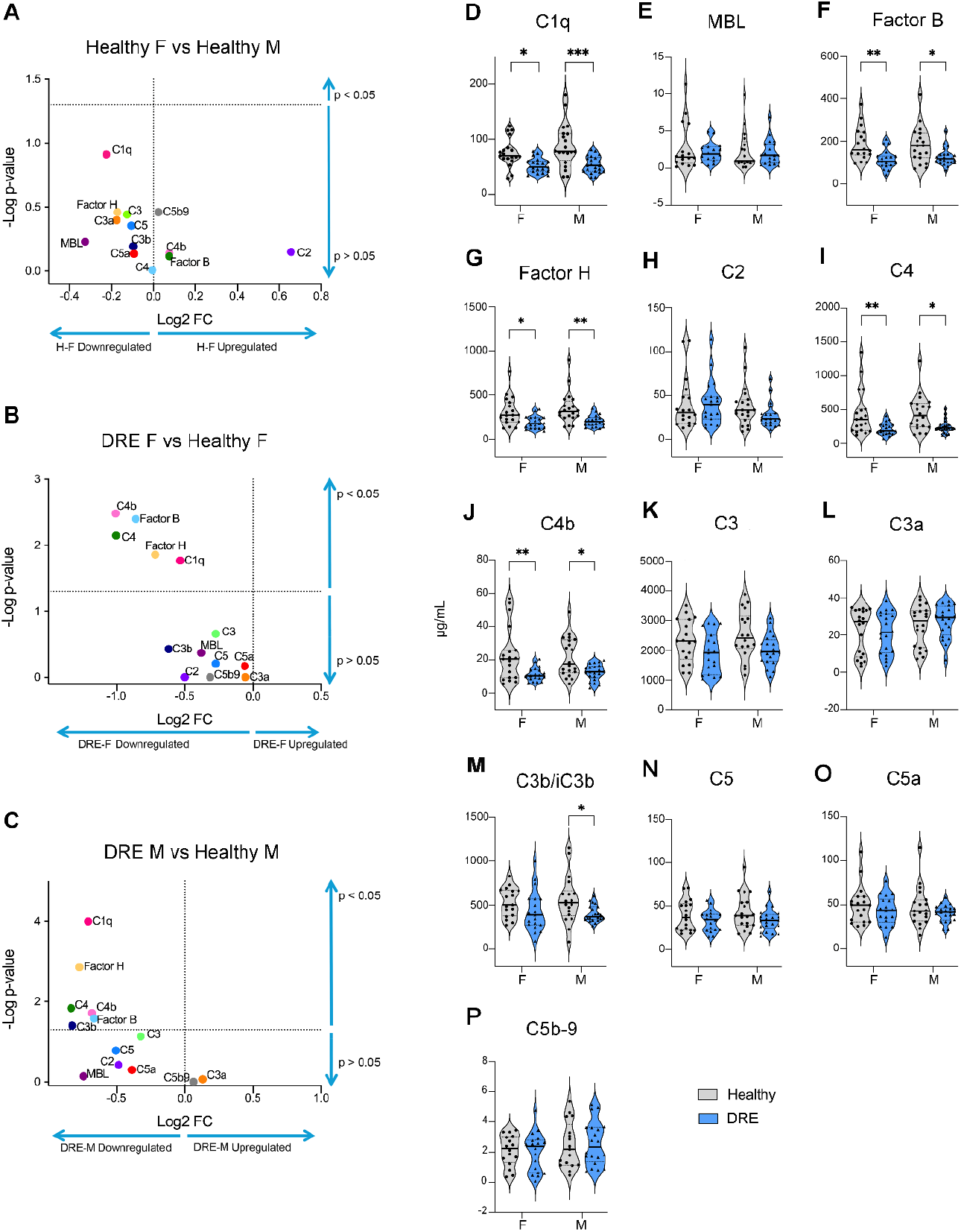
Serum complement component concentrations in male and female patients with drug-resistant epilepsy (DRE) and healthy controls. **A-C**, Volcano plots showing significance (two-way ANOVA) and mean fold change between: (**A**) Healthy females and healthy males (**B**) DRE females and healthy females (**C**) DRE males and healthy males. **D-P**, Serum concentrations of multiple complement molecules: C1q, Mannose-binding Lectin (MBL), Factor B, Factor H, C2, C4, C4b, C3, C3a, C3b/iC3b, C5, C5a, and C5b-9. DRE patients are represented in blue, and healthy controls in gray. Violin plots display individual data points, median, and quartiles. Statistical analysis was performed using two-way ANOVA with categorical variables: condition (DRE or healthy control) and sex (female and male). Significance levels: *p < 0.05, **p < 0.01, ***p < 0.001, ****p < 0.0001.

### Dysregulation of complement pathways in DRE

We further analyzed these molecular differences to elucidate potential functional defects within the complement cascade (Fig. 3, Supplementary Figs. 1-4). The sequential activation of specific components regulates the complement system, necessitating precise coordination to generate active molecules such as C3a, C3b, C5a, and C5b (Fig. 1A) ^33^. In healthy individuals (Figs. 3A and 3C), we observed high positive correlations between classical (C1q to C3b/iC3b), lectin (MBL to C3b/iC3b), and late cleavage pathways (C3b/iC3b to C5a), suggesting that these pathways may be functioning in a coordinated manner to maintain a balanced immune response. However, DRE females showed no correlation between levels of MBL and C3b/iC3b (Fig. 3J), suggesting a targeted impairment within the lectin pathway, while DRE males exhibited a broader lack of correlation between the levels of molecules within both classical (Fig. 3H) and lectin (Fig. 3L) pathways. Although positive C3b/iC3b to C5a correlations were maintained across groups (Fig. 3M-P), a significant positive correlation between C3b/iC3b to C5b-9 (terminal pathway) was only present in healthy males (Supplementary Figures 1-4). These findings suggest a more extensive complement system dysregulation, which could more severely impact immune homeostasis in DRE males.

**Figure 3.**
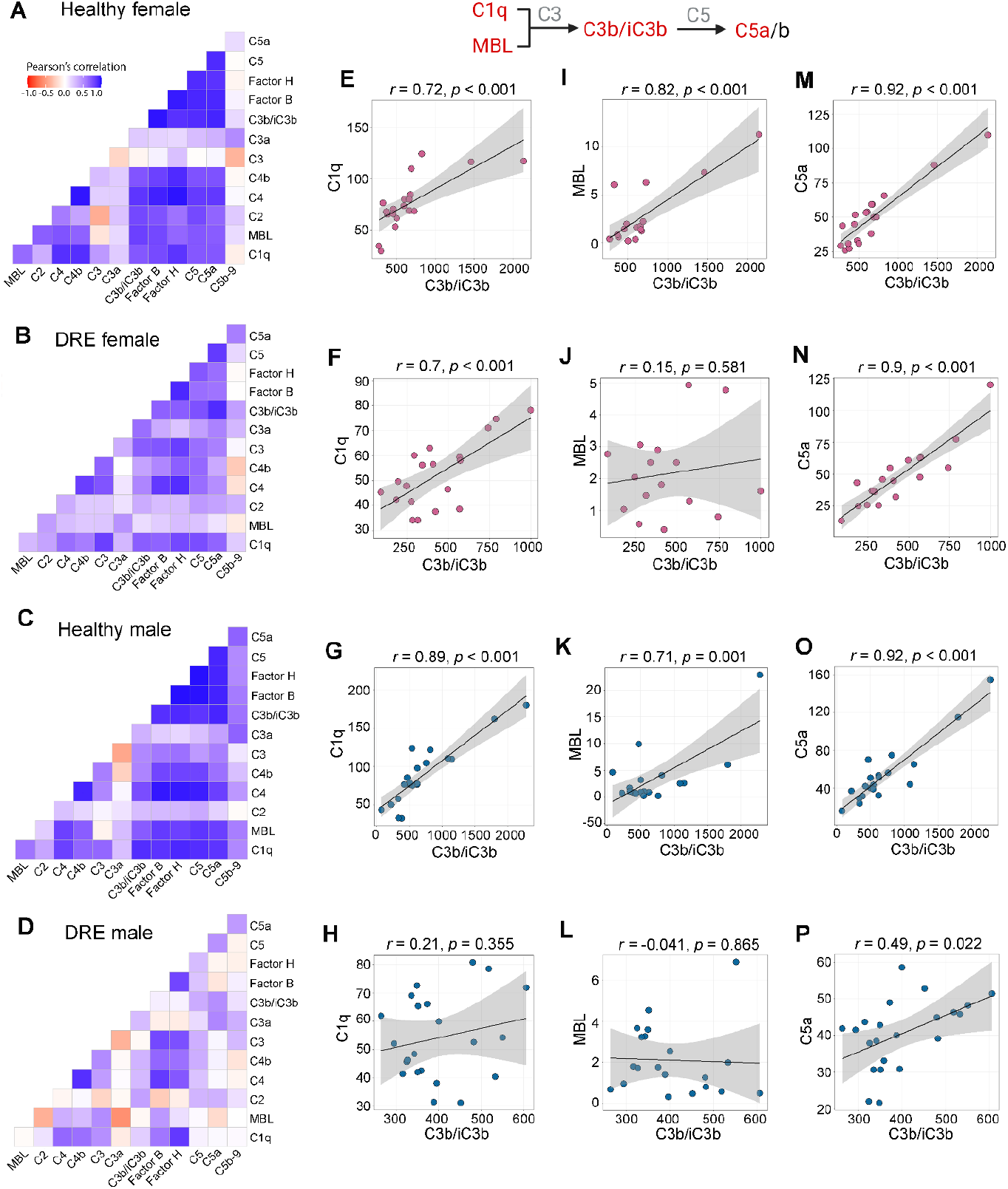
Correlations between complement components. **A-D**, Pearson correlation matrices of all complement components in: (**A**) Healthy females, (**B**) DRE females, (**C**) Healthy males, and (**D**) DRE males. **E-H**, Pearson correlations between C1q and C3b/iC3b in: (**E**) Healthy females, (**F**) DRE females, (**G**) Healthy males, and (**H**) DRE males. **I-L**, Pearson correlations between MBL and C3b/iC3b in: (**I**) Healthy females, (**J**) DRE females, (**K**) Healthy males, and (**L**) DRE males. **M-P**, Pearson correlations between C5a and C3b/iC3b in: (**M**) Healthy females, (**N**) DRE females, (**O**) Healthy males, and (**P**) DRE males. Gray areas in panels **E-P** indicate 95% confidence intervals. Top: Simplified diagram showing the classical (C1q) and lectin (MBL) pathways and downstream cleavage of C3 and C5. Red indicates correlations shown in panels E-P. Supplementary Figures 1-4 show all r and p values for all correlations by group.

### Cytokine serum levels in health and DRE

We investigated the association between the concentrations of complement cascade components and cytokines in serum (Fig. 4-5; Supplementary Figures 1-5). Our findings revealed distinct cytokine profile patterns between healthy individuals and DRE patients (Fig. 4A), as well as between males and females in both healthy (Fig. 4B) and DRE groups (Fig. 4C-D) (Supplementary Tables 1 and 2). First, we observed that the DRE group exhibited significantly lower serum IL-8 levels (p = 0.001) and significantly higher CCL2 (p = 0.03) and CCL5 (p = 0.009) levels compared to the healthy group (Fig. 4A). Additionally, we noted a trend toward significance in the decrease of TNFα levels (p = 0.06) in the DRE group compared to the healthy group. Interestingly, ANOVA revealed significant sex dimorphism in some cytokine levels, with DRE females showing significantly lower levels of TNFα (p < 0.001) and IL-8 (p = 0.007) compared to healthy females (Fig. 4E and 4F), suggesting that these cytokine changes in the DRE group are primarily driven by the DRE female population. In contrast, DRE males showed a trend towards elevated CCL2 (p = 0.125) and CCL5 (p = 0.066) compared to healthy males (Fig. 4G and 4H), although these differences did not reach statistical significance. This trend was not observed in female DRE vs. healthy groups (CCL2, p = 1 ; CCL5, p = 0.261). Notably, bFGF was detected only in the DRE groups, while levels within the healthy group were below the sensitivity limit of the assay kit. We acknowledge that this limitation in detection sensitivity could affect group comparisons.

**Figure 4.**
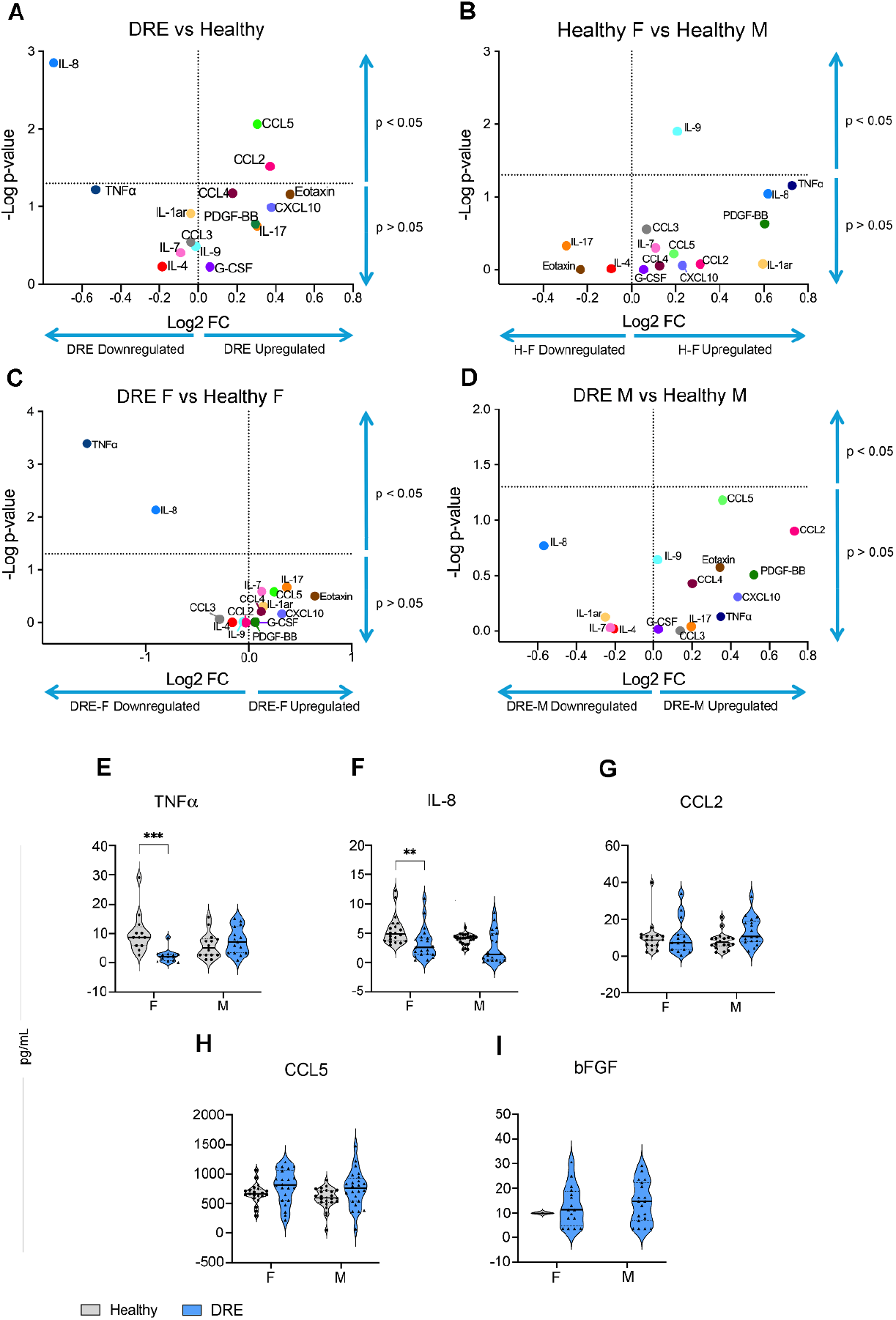
Serum cytokine concentrations in male and female patients with drug-resistant epilepsy (DRE) and healthy controls. **A-D**, Volcano plots showing significance (unpaired *t*-test and two-way ANOVA) and cytokine mean fold change between: (**A**) DRE patients and healthy controls (**B**) Healthy females and healthy males (**C**) DRE females and healthy females (**D**) DRE males and healthy males. **E-I**, Serum concentrations of TNFα, IL-8, CCL2, CCL5, and bFGF in male and female patients. DRE patients are represented in blue, and healthy controls in gray. Violin plots display individual data points, median, and quartiles. Statistical analysis was performed using two-way ANOVA with categorical variables: condition (DRE or healthy control) and sex (female and male). Significance levels: *p < 0.05, **p < 0.01, ***p < 0.001.

Furthermore, we observed marked alterations in the complement and cytokine correlations (Fig. 5). Healthy females (Fig. 5A) and healthy males (Fig. 5C) generally showed more positive correlations, which were less evident in DRE females (Fig. 5B) and DRE males (Fig. 5D). Examples of sex-specific differences include significant positive correlations between C3b/iC3b and IL-17 in females only (Fig. 5E-5H), and negative correlations between C4 and IL-8 in males only (Fig. 5I-5L). Both DRE groups showed significant positive correlations between C3b/iC3b and CCL2 levels that were absent in the corresponding healthy groups (Fig. 5M-5P). The positive correlation between complement and cytokines suggests better immune regulation in healthy individuals. In contrast, altered patterns in DRE patients may indicate maladaptive immune responses.

**Figure 5.**
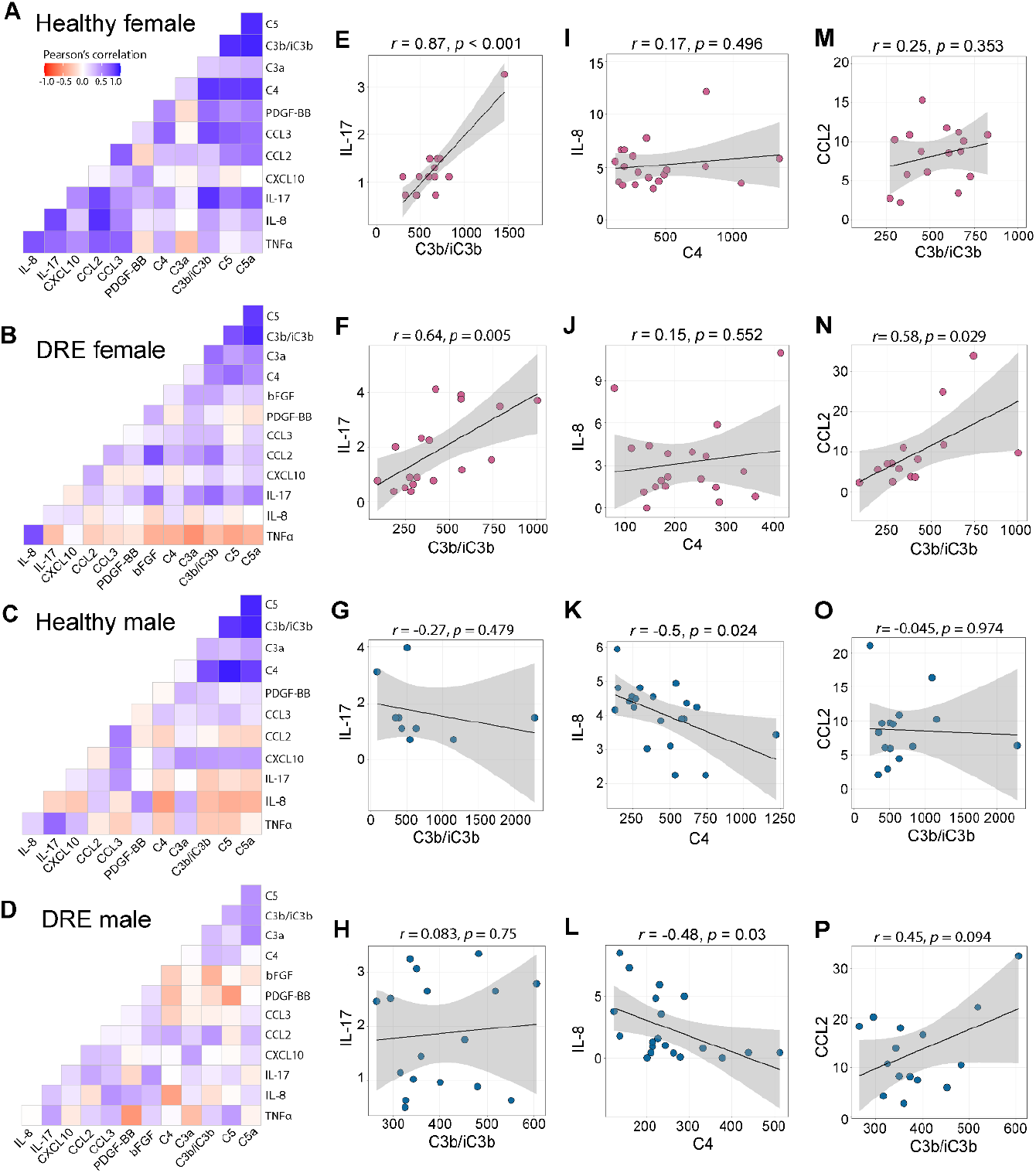
Correlations between serum concentrations of complement components and cytokines. **A-D**, Pearson correlation matrices between complement components and cytokines in: (**A**) Healthy females, (**B**) DRE females, (**C**) Healthy males, and (**D**) DRE males. **E-H**, Pearson correlations between IL-17 and C3b/iC3b in: (**E**) Healthy females, (**F**) DRE females, (**G**) Healthy males, and (**H**) DRE males. **I-L**, Pearson correlations between IL-8 and C4 in: (**I**) Healthy females, (**J**) DRE females, (**K**) Healthy males, and (**L**) DRE males. **M-P**, Pearson correlations between CCL2 and C3b/iC3b in: (**M**) Healthy females, (**N**) DRE females, (**O**) Healthy males, and (**P**) DRE males. Gray areas in panels **E-P** indicate 95% confidence intervals. Supplementary Figures 1-4 show all r and p values for all correlations by group.

### Complement and cytokine correlations with age and clinical outcomes

Finally, we analyzed the associations between complement and cytokine profiles and epilepsy duration, FSIQ scores, and age (Supplementary Figures 1-4). No correlation between epilepsy duration and complement or cytokine levels was found in DRE patients of either sex (Supplementary Figures 1-4). Lower FSIQ scores were correlated with higher levels of C2 and IL-8 in DRE females (Supplementary Fig. 2), while in DRE males, lower FSIQ scores were linked to higher levels of IL-9 (Supplementary Fig. 4). Intriguingly, despite the observed complement deficiency in male DRE patients compared to healthy males, we found a significant positive correlation between serum levels of C1q, C3, and C5 with age (Supplementary Fig. 4), suggesting a potential progressive activation of the complement system with aging in the male DRE group.

## Discussion

Our study presents evidence of complement system deficiency in DRE patients, suggesting potential implications for a seizure-related immune vulnerability that may be exacerbated in a sex-dependent manner. Within the DRE population, we observed a general reduction in the serum levels of multiple complement analytes (C1q, Factor B, C4, C4b, and Factor H) (Fig. 1) along with common increases in bFGF (Fig. 4). Additionally, we found sex-specific differences in both the levels and coordination between complement components and cytokines (Fig. 3-5) in healthy individuals and DRE patients. Our analysis also revealed associations between specific complement and inflammatory markers (C2, IL-8, and IL-9) and FSIQ scores in DRE patients, suggesting their potential as biomarkers for cognitive disability.

Our findings demonstrate lower serum concentrations of complement components such as C1q, C3, and C4 in DRE cases (Figs. 1-2). This evidence aligns with previous research showing alterations in the levels of these molecules in different populations of epilepsy patients ^23, 24^. A study of 157 epilepsy patients found significant decreases in iC3b levels among patients with uncontrolled seizures compared to those with controlled seizures ^23^. Another study involving 37 patients with idiopathic generalized epilepsy reported reduced serum levels of C3 and C4 compared to healthy controls, with a more pronounced decrease in untreated patients ^24^. These findings indicate that patients with epilepsy have a weakened complement system that may potentially render them more susceptible to immune-related comorbid conditions.

A weakened complement system can lead to various health challenges, including compromised innate and adaptive immunity, increased susceptibility to infections, a higher risk of autoimmune diseases, and chronic inflammation ^19-21^. Deficiencies in classical pathway components (C1q, C1, C2, C4) are linked to autoimmune disorders such as systemic lupus erythematosus ^20, 34, 35^. Deficiencies in the MBL, C3, and C5 pathways are also associated with an increased risk of bacterial and viral infections ^19, 21^. Thus, we speculate that reduced levels of complement components in epilepsy patients may increase the risk of developing these immune conditions. Although our literature review did not identify studies linking epilepsy with increased susceptibility to infections or prolonged inflammation, this remains an intriguing area for future research. However, unprovoked seizures have been reported in patients with systemic lupus erythematosus ^20, 36-38^, and C1q knockout mice exhibit neuronal hyperexcitability and epilepsy ^39^, suggesting a plausible mechanistic link between peripheral complement deficiency, seizures, and epilepsy.

Additionally, our findings reveal sex-specific variations in the complement system of DRE patients. Females exhibit a targeted impairment of the lectin pathway, while males show a broader dysregulation of the classical, lectin, and terminal pathways. Our novel findings suggest a more severe complement system dysregulation in males with epilepsy (Fig. 3 and Supplementary Fig. 4). Despite the absence of sex-dependent differences in complement component levels within the healthy groups in this study, previous research has shown significant sex-based variations in both the levels and functional activity of serum complement among healthy individuals ^31, 32^. Research involving 120 healthy Caucasian individuals revealed lower basal serum concentrations of C3, MBL, and C5-C9 components in females compared to males ^32^. Conversely, another study reported higher serum levels of C1q and C3b and enhanced antibody-mediated cell lysis in healthy females, while healthy males showed increased activation of early complement pathway components ^31^. This discrepancy between the current findings and existing literature highlights the complexity of complement system regulation and suggests that factors like sample size, population demographics, or methodological variations (e.g. sensitivity of ELISA kits) may impact the observed outcomes.

The complement system modulates inflammation through its reciprocal relationship with cytokines, creating a feedback loop that initiates and resolves inflammatory responses, thereby maintaining immune homeostasis ^16-18^. Our study revealed common alterations in all DRE cases, including complement deficiencies and detectable bFGF levels (Fig. 4 and Supplementary Figs. 5-6). In DRE females, these changes were accompanied by reduced TNFα and IL-8 levels, while in DRE males, a trend towards elevated CCL2 and CCL5 levels was observed, indicating potential sex dimorphism in their immune signatures. These alterations may affect inflammatory processes. For instance, in DRE females, reduced TNFα may indicate a dampened immune response, potentially affecting inflammation resolution or increasing the risk of chronic inflammation, as lower levels of TNFα are also linked to increases in IL-6 ^40^.

Furthermore, reduced IL-8, which is crucial for neutrophil recruitment, infection resolution, wound healing, and angiogenesis ^41^, could affect responses to infections and tissue damage. In contrast, the trend towards elevated CCL2 and CCL5, pro-inflammatory chemokines that recruit monocytes and lymphocytes ^42^, suggests a potential shift in inflammatory state in DRE males. The presence of detectable bFGF in DRE cases, but not in healthy controls, suggests a potential compensatory mechanism to enhance tissue repair and angiogenesis ^43^. While these findings provide interesting insights into the immune profiles of DRE patients, a more comprehensive understanding of cytokine and chemokine profiles is essential for drawing targeted conclusions, particularly given the observed trends and sex-specific differences.

A growing body of evidence in human and animal studies demonstrates sex-based disparities in cytokine production and immune responses in both health and disease conditions ^26-30^, an aspect understudied in epilepsy. For example, studies of healthy human subjects revealed that males have higher plasma levels of TNFα, IL-1β, and IL-6 ^27^ and a stronger monocyte-derived pro-inflammatory cytokine production in response to *in vitro* lipopolysaccharide challenge of blood samples than females ^29^. In association with critical trauma events, males exhibit higher circulating concentrations of cytokines like TNFα and IFN-γ ^28^. In the context of Alzheimer’s disease, evaluation of inflammatory profiles in peripheral blood leukocytes revealed higher levels of TNFα and IL-1β in males than females ^30^. These findings collectively indicate sex-based differences in cytokine levels, with males generally displaying enhanced pro-inflammatory profiles compared to females in both healthy and injury or disease conditions. This pattern aligns with our findings, showing that DRE males exhibit elevated CCL2 and CCL5 levels. The alterations in these chemokine levels may be linked to the more extensive complement system dysregulation observed in DRE males. However, further research is necessary to confirm this potential connection.

Several limitations warrant consideration in future research. It remains unclear whether the observed weakened complement system and the altered landscape of cytokine/chemokine profiles cause the seizures or are a result of them, and whether they are beneficial or harmful in epilepsy. Moreover, these alterations could also be attributed to the use of ASMs, as these medications can affect immune function ^44^. Some ASMs, such as phenytoin, carbamazepine, and valproate, have immunosuppressive effects ^44^, which may increase the risk of infection ^45^. Additionally, immune system function can be influenced by various factors, including sex hormones ^46^ and age ^47^. The timing of blood sampling is also critical, as interictal and ictal phases could present different blood biomarker profiles. These limitations underscore the need for comprehensive, controlled studies that account for ASMs and other medication effects, health comorbidities, age, and seizure timing. Such studies would further elucidate the complex relationship between the complement system, immune responses, and uncontrolled versus controlled seizures.

The concept of a vulnerable immune profile leading to maladaptive inflammatory responses in DRE cases aligns with previous research showing changes in CD4+ T cell abundance, a key component of the immune system, in the blood of epilepsy patients. These changes vary across different types of epilepsy with decreased proportions of CD4+ T cells reported in temporal lobe epilepsy ^11^ and focal epilepsy of unknown cause ^12^, and increased abundance found in DRE cases ^13^. Despite these variations, all these epilepsy types often exhibit elevated inflammatory cytokines ^11-14^. However, the relationship between peripheral immune cells, cytokines, the complement cascade, and their impact on immune responses or epilepsy outcomes likely depends on a broader set of immune regulators, which may also vary by sex. Taken together, our findings emphasize the importance of sex in immune signatures, an aspect often overlooked in clinical and pre-clinical epilepsy research. The ultimate goal is that our findings help develop clinical management strategies accounting for the individual variability in immunological trajectories and to improve outcomes for patients with DRE.

## Acknowledgments

This research was supported by: National Institute of Neurological Disorders and Stroke, Grant/Award Number, NS096234 (ALB); Children’s Brain Diseases Foundation (ALB), Laboratory startup funds SMU (ALB). ANR EpiNeurAge and EpiCatcher to NM. We thank IU Health ECRO Biorepository for providing us with all the serum samples utilized in this study.

## Author Contributions

NPH: Conceptualization, Investigation, Methodology, Validation, Data curation, Formal analysis, Writing – original Draft, Writing – review and editing. YL: Investigation, Methodology, Data curation, Formal analysis, Writing – review and editing. MM: Formal analysis, Writing – review and editing. NPP: Writing – review and editing. NM: Writing – review and editing. AB: Conceptualization, Data curation, Formal analysis, Funding acquisition, Resources, Supervision, Writing – original draft, Writing – review and editing.

## Data Availability

This study presents a comprehensive dataset sufficient for thorough evaluation of the experimental outcomes. The data supporting these findings are available in the main article and accompanying supplementary material, with additional materials available from the corresponding author upon written request.

## References

1. Asadi-Pooya AA, Stewart GR, Abrams DJ, Sharan A. Prevalence and Incidence of Drug-Resistant Mesial Temporal Lobe Epilepsy in the United States. World Neurosurg. 2017 Mar;99:662–6.

2. van Vliet EA, Marchi N. Neurovascular unit dysfunction as a mechanism of seizures and epilepsy during aging. Epilepsia. 2022 Jun;63(6):1297–313.

3. Hanin A, Cespedes J, Dorgham K, et al. Cytokines in New-Onset Refractory Status Epilepticus Predict Outcomes. Ann Neurol. 2023 Jul;94(1):75–90.

4. Villasana-Salazar B, Vezzani A. Neuroinflammation microenvironment sharpens seizure circuit. Neurobiol Dis. 2023 Mar;178:106027.

5. Janigro D, Bailey DM, Lehmann S, et al. Peripheral Blood and Salivary Biomarkers of Blood-Brain Barrier Permeability and Neuronal Damage: Clinical and Applied Concepts. Front Neurol. 2020;11:577312.

6. Hosseini S, Mofrad AME, Mokarian P, et al. Neutrophil to Lymphocyte Ratio in Epilepsy: A Systematic Review. Mediators Inflamm. 2022;2022:4973996.

7. Lim HK, Bae S, Han K, et al. Seizure-induced neutrophil adhesion in brain capillaries leads to a decrease in postictal cerebral blood flow. iScience. 2023 May 19;26(5):106655.

8. Stredny C, Rotenberg A, Leviton A, Loddenkemper T. Systemic inflammation as a biomarker of seizure propensity and a target for treatment to reduce seizure propensity. Epilepsia Open. 2023 Mar;8(1):221–34.

9. Marchi N, Granata T, Janigro D. Inflammatory pathways of seizure disorders. Trends Neurosci. 2014 Feb;37(2):55–65.

10. Shin HR, Chu K, Lee WJ, et al. Neuropsychiatric symptoms and seizure related with serum cytokine in epilepsy patients. Sci Rep. 2022 May 3;12(1):7138.

11. Bauer S, Koller M, Cepok S, et al. NK and CD4+ T cell changes in blood after seizures in temporal lobe epilepsy. Exp Neurol. 2008 Jun;211(2):370–7.

12. Sanli E, Sirin NG, Kucukali CI, et al. Peripheral blood regulatory B and T cells are decreased in patients with focal epilepsy. J Neuroimmunol. 2024 Feb 15;387:578287.

13. Ouedraogo O, Rebillard RM, Jamann H, et al. Increased frequency of proinflammatory CD4 T cells and pathological levels of serum neurofilament light chain in adult drug-resistant epilepsy. Epilepsia. 2021 Jan;62(1):176–89.

14. Wang J, Wu Y, Chen J, et al. Th1/Th2 Imbalance in Peripheral Blood Echoes Microglia State Dynamics in CNS During TLE Progression. Adv Sci (Weinh). 2024 Aug 13:e2405346.

15. Gao F, Gao Y, Zhang SJ, et al. Alteration of plasma cytokines in patients with active epilepsy. Acta Neurol Scand. 2017 Jun;135(6):663–9.

16. Lo MW, Woodruff TM. Complement: Bridging the innate and adaptive immune systems in sterile inflammation. J Leukoc Biol. 2020 Jul;108(1):339–51.

17. Mastellos DC, Hajishengallis G, Lambris JD. A guide to complement biology, pathology and therapeutic opportunity. Nat Rev Immunol. 2024 Feb;24(2):118–41.

18. Schartz ND, Tenner AJ. The good, the bad, and the opportunities of the complement system in neurodegenerative disease. J Neuroinflammation. 2020 Nov 25;17(1):354.

19. Schroder-Braunstein J, Kirschfink M. Complement deficiencies and dysregulation: Pathophysiological consequences, modern analysis, and clinical management. Mol Immunol. 2019 Oct;114:299–311.

20. van Schaarenburg RA, Magro-Checa C, Bakker JA, et al. C1q Deficiency and Neuropsychiatric Systemic Lupus Erythematosus. Front Immunol. 2016;7:647.

21. Bernacchia A, Ginaca A, Rotondo S, Tejada MP, Di Giovanni D. Case Report: C3 deficiency in two siblings. Front Pediatr. 2024;12:1424380.

22. Petrisko TJ, Gomez-Arboledas A, Tenner AJ. Complement as a powerful “influencer” in the brain during development, adulthood and neurological disorders. Adv Immunol. 2021;152:157–222.

23. Kopczynska M, Zelek WM, Vespa S, et al. Complement system biomarkers in epilepsy. Seizure. 2018 Aug;60:1–7.

24. Liguori C, Romigi A, Izzi F, et al. Complement system dysregulation in patients affected by Idiopathic Generalized Epilepsy and the effect of antiepileptic treatment. Epilepsy Res. 2017 Nov;137:107–11.

25. Ma M, Cheng Y, Hou X, et al. Serum biomarkers in patients with drug-resistant epilepsy: a proteomics-based analysis. Front Neurol. 2024;15:1383023.

26. Osborne BF, Turano A, Schwarz JM. Sex Differences in the Neuroimmune System. Curr Opin Behav Sci. 2018 Oct;23:118–23.

27. Bernardi S, Toffoli B, Tonon F, et al. Sex Differences in Proatherogenic Cytokine Levels. Int J Mol Sci. 2020 May 29;21(11).

28. Guidry CA, Swenson BR, Davies SW, et al. Sex- and diagnosis-dependent differences in mortality and admission cytokine levels among patients admitted for intensive care. Crit Care Med. 2014 May;42(5):1110–20.

29. Beenakker KGM, Westendorp RGJ, de Craen AJM, et al. Men Have a Stronger Monocyte-Derived Cytokine Production Response upon Stimulation with the Gram-Negative Stimulus Lipopolysaccharide than Women: A Pooled Analysis Including 15 Study Populations. J Innate Immun. 2020;12(2):142–53.

30. Sochocka M, Ochnik M, Sobczynski M, Orzechowska B, Leszek J. Sex Differences in Innate Immune Response of Peripheral Blood Leukocytes of Alzheimer’s Disease Patients. Arch Immunol Ther Exp (Warsz). 2022 Jun 16;70(1):16.

31. Kelkar NS, Goldberg BS, Dufloo J, et al. Sex- and species-associated differences in complement-mediated immunity in humans and rhesus macaques. mBio. 2024 Mar 13;15(3):e0028224.

32. Gaya da Costa M, Poppelaars F, van Kooten C, et al. Age and Sex-Associated Changes of Complement Activity and Complement Levels in a Healthy Caucasian Population. Front Immunol. 2018;9:2664.

33. Barnum SR. Complement: A primer for the coming therapeutic revolution. Pharmacol Ther. 2017 Apr;172:63–72.

34. Kleer JS, Skattum L, Dubler D, et al. Complement C1s deficiency in a male Caucasian patient with systemic lupus erythematosus: a case report. Front Immunol. 2023;14:1257525.

35. Schejbel L, Skattum L, Hagelberg S, et al. Molecular basis of hereditary C1q deficiency--revisited: identification of several novel disease-causing mutations. Genes Immun. 2011 Dec;12(8):626–34.

36. Mehta P, Norsworthy PJ, Hall AE, et al. SLE with C1q deficiency treated with fresh frozen plasma: a 10-year experience. Rheumatology (Oxford). 2010 Apr;49(4):823–4.

37. Hannema AJ, Kluin-Nelemans JC, Hack CE, Eerenberg-Belmer AJ, Mallee C, van Helden HP. SLE like syndrome and functional deficiency of C1q in members of a large family. Clin Exp Immunol. 1984 Jan;55(1):106–14.

38. Rodriguez-Hernandez A, Ortiz-Orendain J, Alvarez-Palazuelos LE, Gonzalez-Lopez L, Gamez-Nava JI, Zavala-Cerna MG. Seizures in systemic lupus erythematosus: A scoping review. Seizure. 2021 Mar;86:161–7.

39. Chu Y, Jin X, Parada I, et al. Enhanced synaptic connectivity and epilepsy in C1q knockout mice. Proc Natl Acad Sci U S A. 2010 Apr 27;107(17):7975–80.

40. Tuazon Kels MJ, Ng E, Al Rumaih Z, et al. TNF deficiency dysregulates inflammatory cytokine production, leading to lung pathology and death during respiratory poxvirus infection. Proc Natl Acad Sci U S A. 2020 Jul 7;117(27):15935–46.

41. Matsushima K, Yang D, Oppenheim JJ. Interleukin-8: An evolving chemokine. Cytokine. 2022 May;153:155828.

42. Gschwandtner M, Derler R, Midwood KS. More Than Just Attractive: How CCL2 Influences Myeloid Cell Behavior Beyond Chemotaxis. Front Immunol. 2019;10:2759.

43. Xie Y, Su N, Yang J, et al. FGF/FGFR signaling in health and disease. Signal Transduct Target Ther. 2020 Sep 2;5(1):181.

44. Beghi E, Shorvon S. Antiepileptic drugs and the immune system. Epilepsia. 2011 May;52 Suppl 3:40–4.

45. Zaccara G, Giovannelli F, Giorgi FS, Franco V, Gasparini S, Tacconi FM. Do antiepileptic drugs increase the risk of infectious diseases? A meta-analysis of placebo-controlled studies. Br J Clin Pharmacol. 2017 Sep;83(9):1873–9.

46. Sciarra F, Campolo F, Franceschini E, Carlomagno F, Venneri MA. Gender-Specific Impact of Sex Hormones on the Immune System. Int J Mol Sci. 2023 Mar 27;24(7).

47. Lewis ED, Wu D, Meydani SN. Age-associated alterations in immune function and inflammation. Prog Neuropsychopharmacol Biol Psychiatry. 2022 Aug 30;118:110576.

