## Supplementary data for "Sex-Specific Complement and Cytokine Imbalances in Drug-Resistant Epilepsy: Biomarkers of Immune Vulnerability"

Running Title: Sex-Specific Complement Dysregulation in Refractory Epilepsy

Authors: Nicole Pinzon-Hoyos, BS,[1] Yibo Li, MS,[1] Monnie McGee, PhD,[2] Nicholas P. Poolos, MD PhD,[3] Nicola Marchi, PhD, [4] Amy L. Brewster, PhD[* 1]

[1]Department of Biological Sciences, Southern Methodist University, Dallas, TX, USA. [2]Department of Statistics and Data Science, Southern Methodist University, Dallas, TX, USA.

[3]Department of Neurology and Regional Epilepsy Center, University of Washington, Seattle, WA, USA.

[4]Institut de Génomique Fonctionnelle, University of Montpellier, CNRS, INSERM; Cerebrovascular and Glia Research, Montpellier, France.

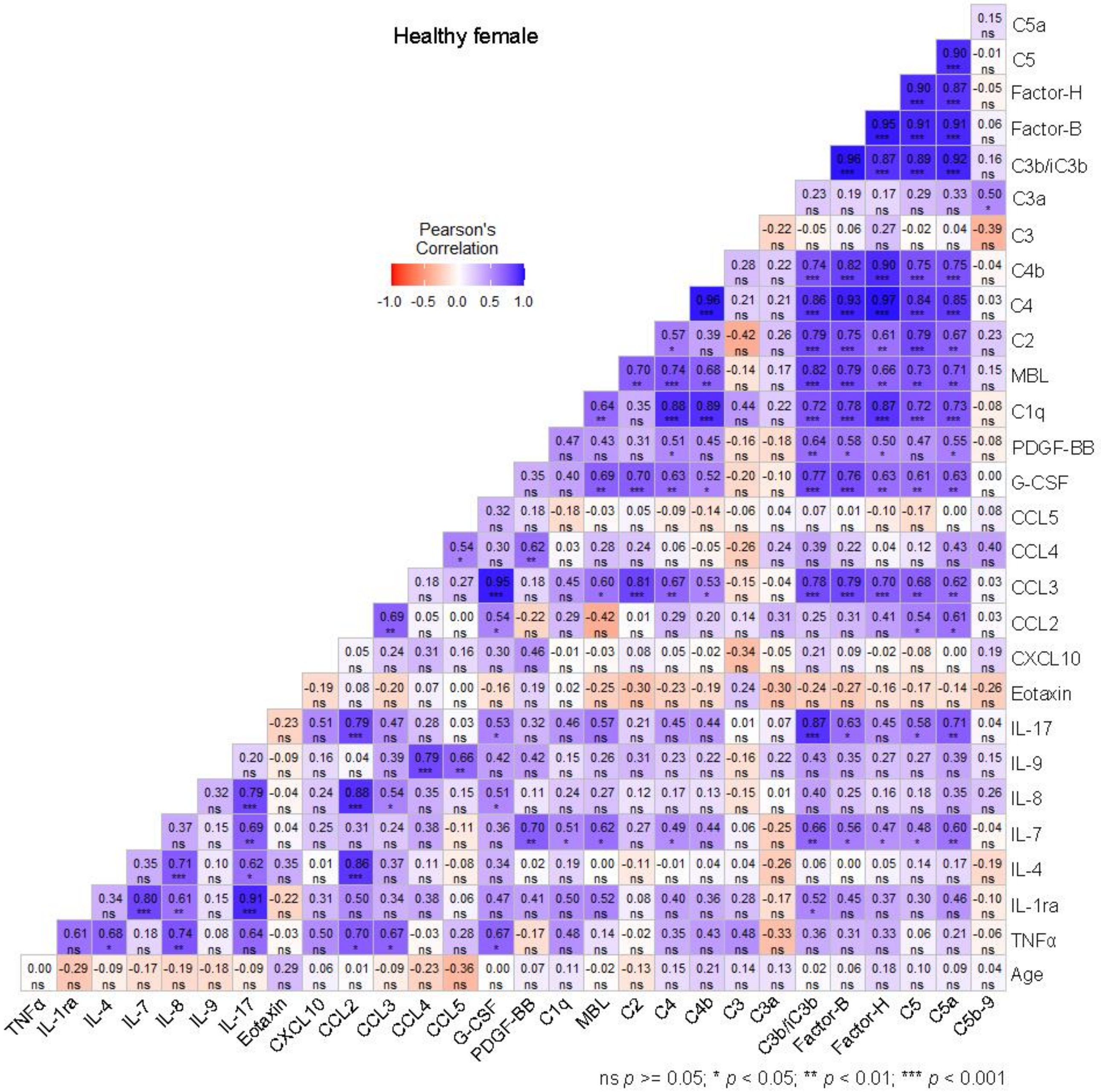


**Supplementary Figure 1.** Pearson correlations between serum concentrations of complement components, cytokines/chemokines, epilepsy duration, FSIQ scores, and age at collection in healthy females. r values are shown for each comparison. *p < 0.05, **p < 0.01, ***p < 0.001. Not significant (ns) p > 0.05.

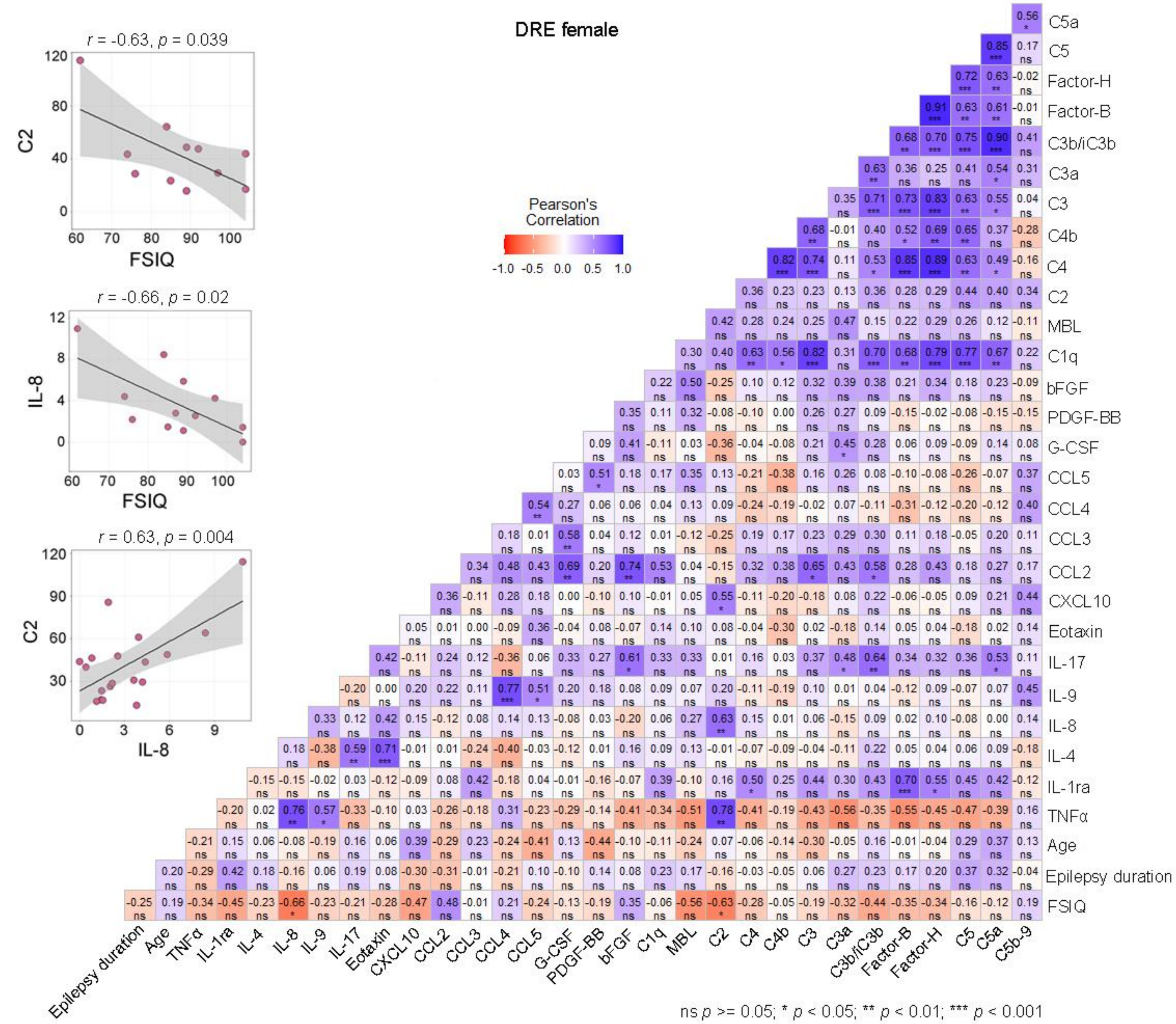


**Supplementary Figure 2.** Pearson correlations between serum concentrations of complement components, cytokines/chemokines, epilepsy duration, FSIQ scores, and age at collection in DRE females. Pearson correlation graphs between C2, IL-8, and FSIQ are shown. r values are shown for each comparison. *p < 0.05, **p < 0.01, ***p < 0.001. Not significant (ns) p > 0.05.

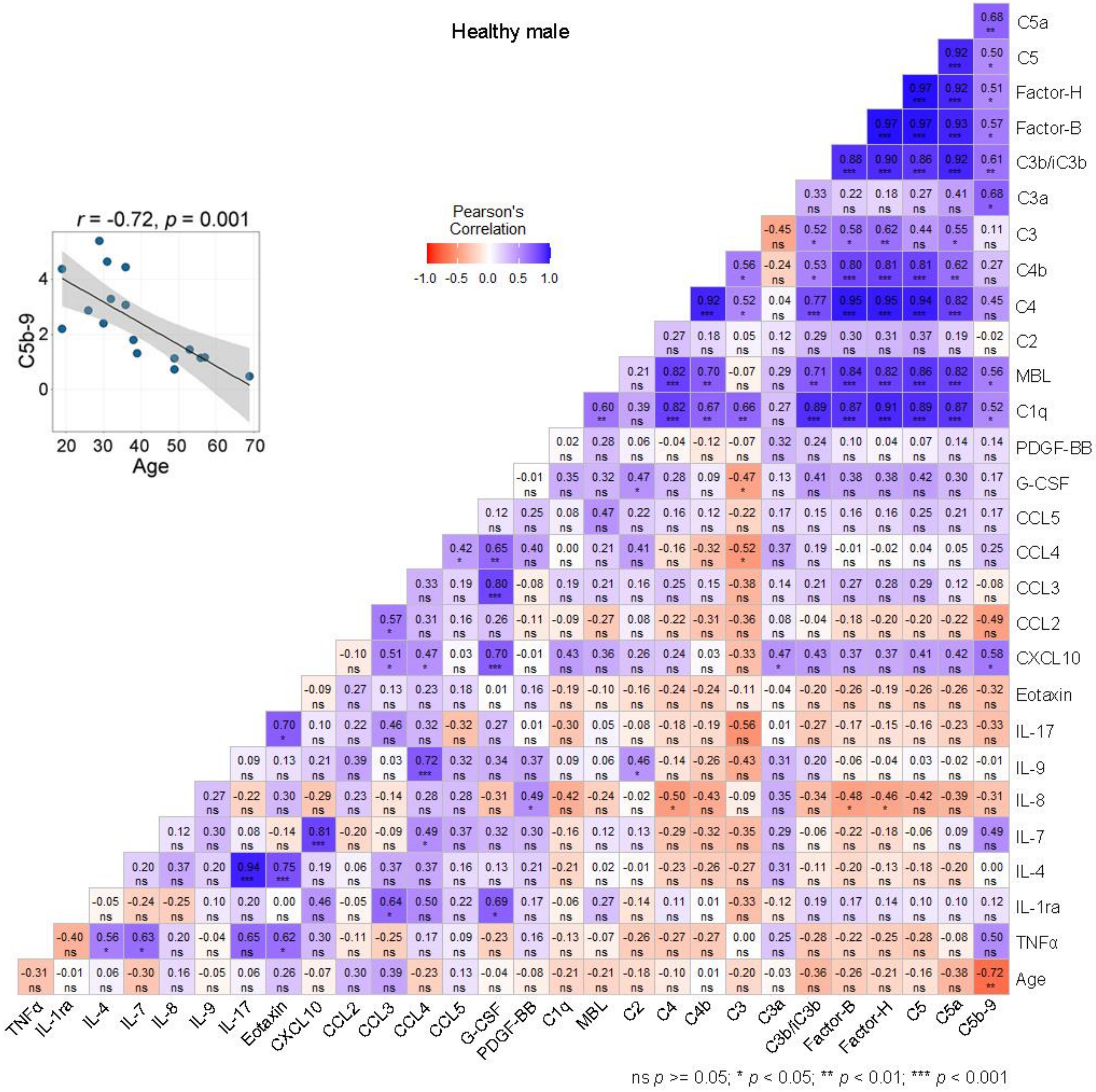


**Supplementary Figure 3.** Pearson correlations between serum concentrations of complement components, cytokines/chemokines, epilepsy duration, FSIQ scores, and age at collection in healthy males. Pearson correlation graph between C5b-9 and age is shown. r values are shown for each comparison. *p < 0.05, **p < 0.01, ***p < 0.001. Not significant (ns) p > 0.05.

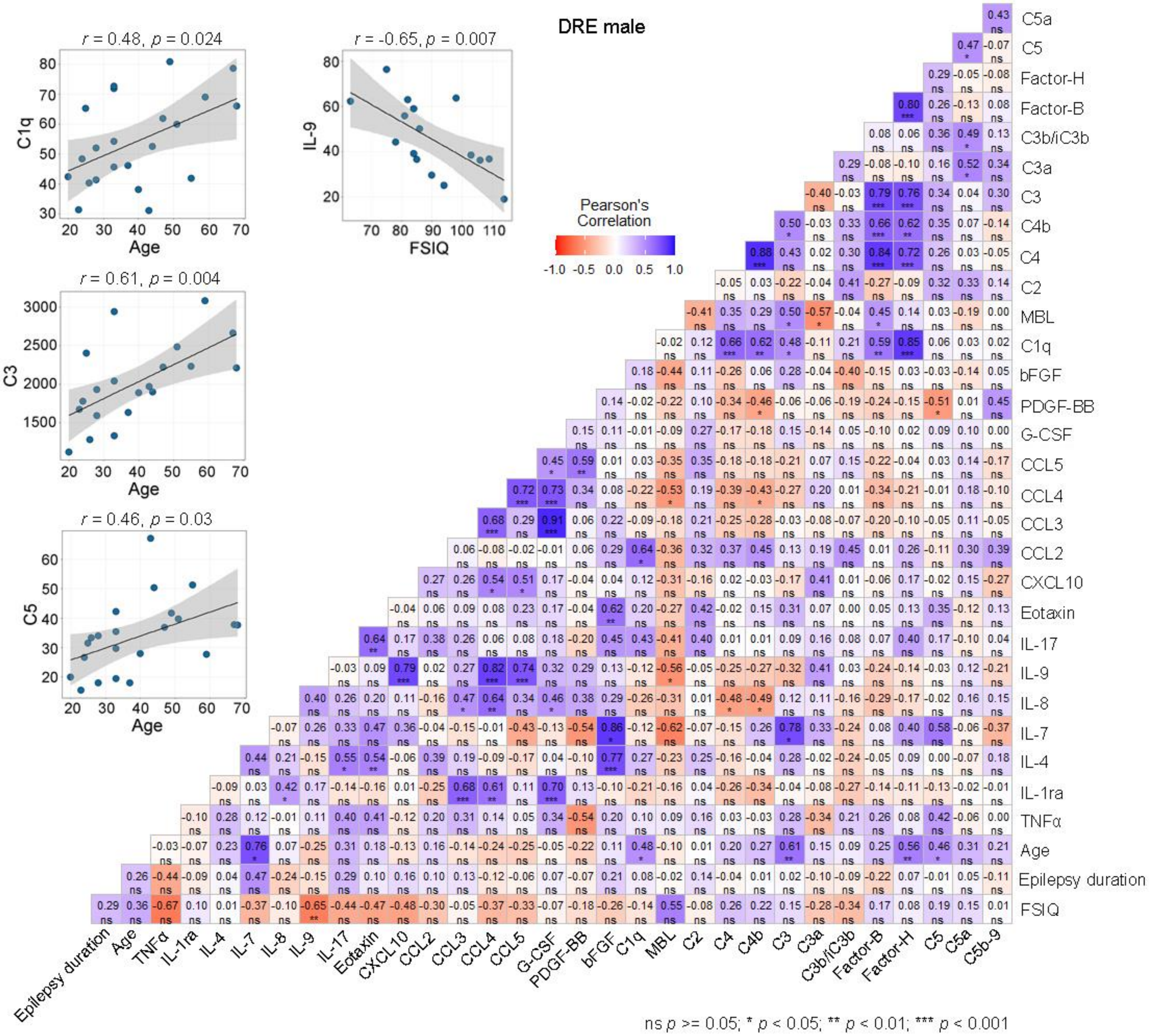


**Supplementary Figure 4.** Pearson correlations between serum concentrations of complement components, cytokines/chemokines, epilepsy duration, FSIQ scores, and age at collection in DRE males. Pearson correlation graphs between C1q, C3, and C5 with age, and IL-9 with FSIQ are shown. r values are shown for each comparison. *p < 0.05, **p < 0.01, ***p < 0.001. Not significant (ns) p > 0.05.

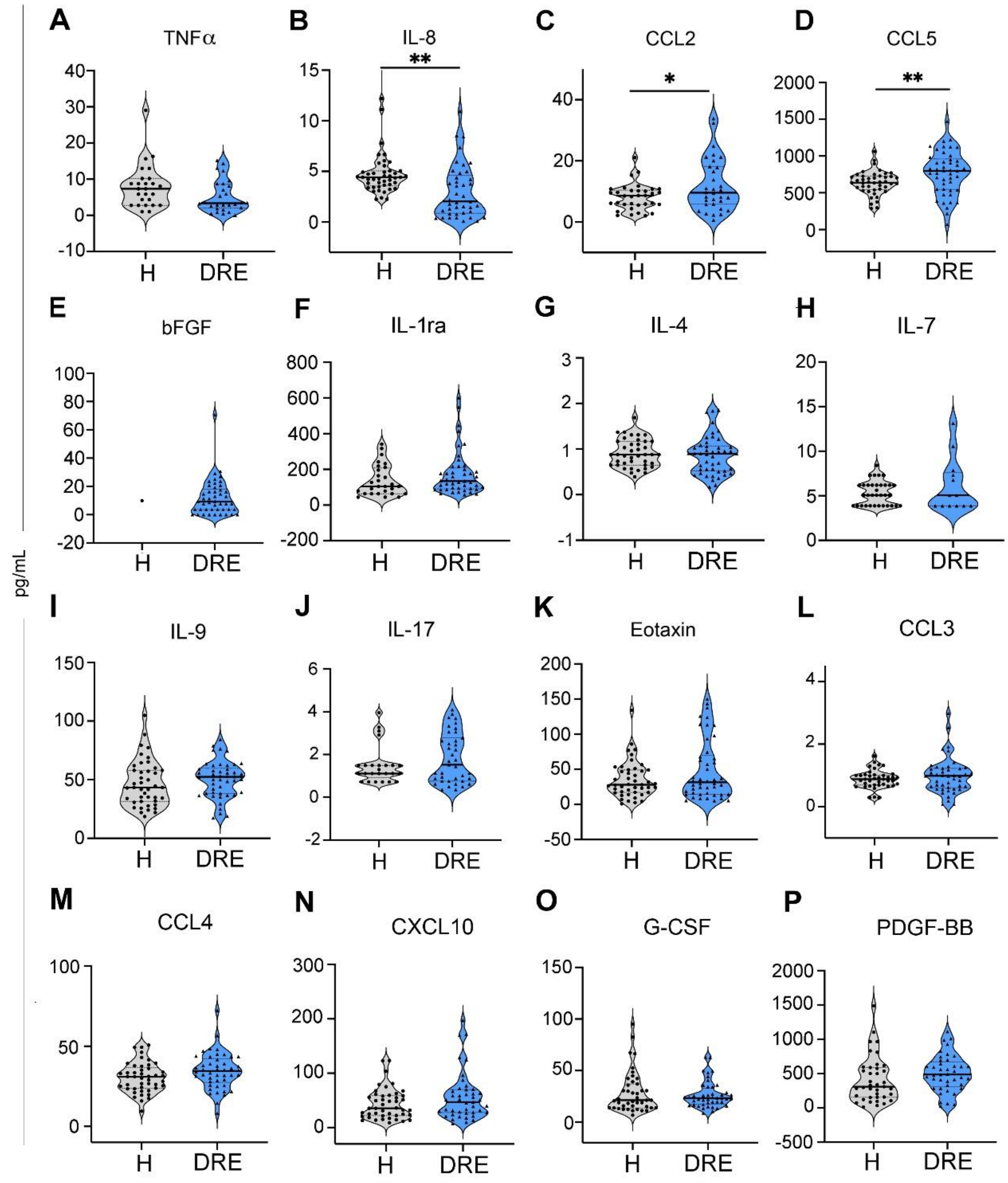


**Supplementary Figure 5.** Serum cytokine concentrations in patients with drug-resistant epilepsy (DRE) and healthy controls. **A-P,** Serum levels of: TNFα, IL-8, CCL2, CCL5, bFGF, IL-1ra, IL-4, IL-7, IL-9, IL-17, Eotaxin, CCL3, CCL4, CXCL10, G-CSF, and PDGF-BB. Statistical analysis: Unpaired t-test. DRE patients are represented in blue, and healthy controls in gray. Violin plots display individual data points, median, and quartiles. Significance levels: $*p < 0.05$, $**p < 0.01$.

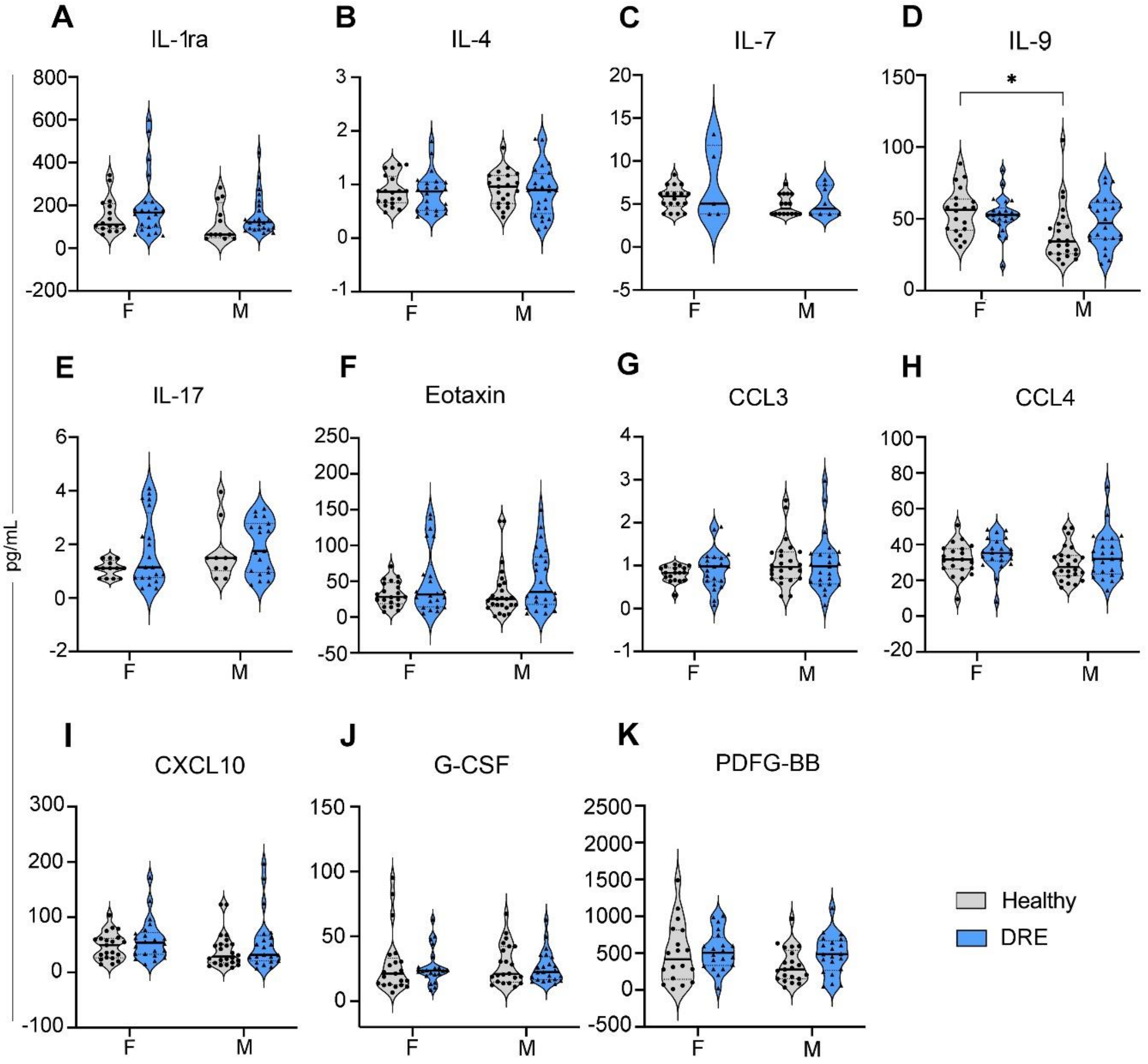


**Supplementary Figure 6.** Serum cytokine concentrations in female and male patients with drug-resistant epilepsy (DRE) and healthy controls. **A-K,** Serum concentrations of IL-1ra, IL-4, IL-7, IL-9, IL-17, Eotaxin, CCL3, CCL4, CXCL10, G-CSF, and PDGF-BB in healthy and DRE groups by sex. Statistical analysis: Two-way ANOVA with categorical variables: condition (DRE or healthy control) and sex (female and male). DRE patients are represented in blue, and healthy controls in gray. Violin plots display individual data points, median, and quartiles. Significance levels: *p < 0.05.

| Analyte | p value |
|---|---|
| C1q | <0.0001 |
| MBL | 0.417 |
| Factor B | <0.001 |
| Factor H | <0.0001 |
| C2 | 0.35 |
| C4 | <0.001 |
| C4b | <0.0001 |
| C3 | 0.006 |
| C3a | 0.744 |
| C3b/iC3b | 0.006 |
| C5 | 0.038 |
| C5a | 0.104 |
| C5b-9 | 0.968 |
| TNFα | 0.061 |
| IL-8 | 0.001 |
| CCL2 | 0.03 |
| CCL5 | 0.009 |
| bFGF | NA |
| IL-1ra | 0.122 |
| IL-4 | 0.591 |
| IL-7 | 0.393 |
| IL-9 | 0.324 |
| IL-17 | 0.178 |
| Eotaxin | 0.069 |
| CCL3 | 0.287 |
| CCL4 | 0.067 |
| CXCL10 | 0.102 |
| G-CSF | 0.595 |
| PDGF-BB | 0.168 |

**Supplementary Table 1.** P values (unpaired t-test with Welch's correction) of serum complement components and cytokine concentrations in DRE and healthy groups.

| ANOVA | | | | Šídák's multiple comparisons test p values | | |
|---|---|---|---|---|---|---|
| Analyte | Interaction | Sex | Condition | H-F vs H-M | H-F vs DRE-F | H-M vs DRE-M |
| C1q | F (1, 76) = 0.9203, p = 0.340 | F (1, 76) = 1.626, p = 0.206 | F (1, 76) = 25.77, p < 0.0001 | 0.324 | 0.017 | <0.001 |
| MBL | F (1, 65) = 0.1145, p = 0.736 | F (1, 65) = 0.1948, p = 0.660 | F (1, 65) = 0.6920, p = 0.409 | 0.931 | 0.811 | 0.978 |
| Factor B | F (1, 74) = 0.3178, p = 0.575 | F (1, 74) = 0.01633, p = 0.899 | F (1, 74) = 18.24, p < 0.0001 | 0.987 | 0.004 | 0.026 |
| Factor H | F (1, 76) = 0.1648, p = 0.686 | F (1, 76) = 0.9053, p = 0.344 | F (1, 76) = 21.43, p < 0.0001 | 0.72 | 0.014 | 0.001 |
| C2 | F (1, 75) = 1.296, p = 0.259 | F (1, 75) = 2.824, p = 0.097 | F (1, 75) = 0.7615, p = 0.386 | 0.975 | 0.997 | 0.377 |
| C4 | F (1, 76) = 0.07756, p = 0.781 | F (1, 76) = 0.09642, p = 0.757 | F (1, 76) = 18.28, p < 0.0001 | 1 | 0.007 | 0.015 |
| C4b | F (1, 75) = 0.2524, p = 0.617 | F (1, 75) = 1.856e-006, p = 1 | F (1, 75) = 19.29, p < 0.0001 | 0.979 | 0.003 | 0.019 |
| C3 | F (1, 71) = 0.1162, p = 0.734 | F (1, 71) = 0.9695, p = 0.328 | F (1, 71) = 8.260, p = 0.005 | 0.739 | 0.22 | 0.074 |
| C3a | F (1, 76) = 0.4477, p = 0.506 | F (1, 76) = 3.485, p = 0.066 | F (1, 76) = 0.1101, p = 0.741 | 0.784 | 0.993 | 0.86 |
| C3b/iC3b | F (1, 72) = 0.4631, p = 0.498 | F (1, 72) = 5.566e-005, p = 0.994 | F (1, 72) = 7.968, p = 0.006 | 0.954 | 0.372 | 0.039 |
| C5 | F (1, 73) = 0.2631, p = 0.610 | F (1, 73) = 0.3614, p = 0.550 | F (1, 73) = 4.490, p = 0.038 | 0.825 | 0.618 | 0.166 |
| C5a | F (1, 74) = 0.02046, p = 0.887 | F (1, 74) = 0.4062, p = 0.526 | F (1, 74) = 2.600, p = 0.111 | 0.98 | 0.677 | 0.501 |
| C5b-9 | F (1, 67) = 0.1248, p = 0.725 | F (1, 67) = 2.003, p = 0.162 | F (1, 67) = 9.151e-005, p = 0.992 | 0.851 | 0.993 | 0.991 |
| TNFα | F (1, 50) = 13.44, p < 0.001 | F (1, 50) = 0.09530, p = 0.759 | F (1, 50) = 5.840, p = 0.019 | 0.07 | <0.001 | 0.742 |
| IL-8 | F (1, 81) = 0.8501, p = 0.359 | F (1, 81) = 4.528, p = 0.036 | F (1, 81) = 12.77, p < 0.001 | 0.091 | 0.007 | 0.17 |
| CCL2 | F (1, 63) = 1.947, p = 0.168 | F (1, 63) = 0.1267, p = 0.723 | F (1, 63) = 2.097, p = 0.153 | 0.838 | 1 | 0.125 |
| CCL5 | F (1, 87) = 0.1392, p = 0.710 | F (1, 87) = 1.477, p = 0.228 | F (1, 87) = 7.950, p = 0.006 | 0.607 | 0.261 | 0.066 |
| bFGF | N/A | N/A | N/A | N/A | N/A | N/A |
| IL-1ra | F (1, 69) = 0.04170, p = 0.839 | F (1, 69) = 1.963, p = 0.166 | F (1, 69) = 2.353, p = 0.130 | 0.834 | 0.496 | 0.749 |
| IL-4 | F (1, 83) = 0.008677, p = 0.926 | F (1, 83) = 0.1971, p = 0.658 | F (1, 83) = 0.2749, p = 0.602 | 0.976 | 0.99 | 0.96 |
| IL-7 | F (1, 43) = 0.9141, p = 0.344 | F (1, 43) = 5.195, p = 0.028 | F (1, 43) = 2.694, p = 0.108 | 0.501 | 0.257 | 0.937 |
| IL-9 | F (1, 85) = 2.228, p = 0.139 | F (1, 85) = 7.528, p = 0.007 | F (1, 85) = 0.8476, p = 0.360 | 0.013 | 0.972 | 0.228 |
| IL-17 | F (1, 58) = 0.5857, p = 0.447 | F (1, 58) = 1.795, p = 0.186 | F (1, 58) = 2.749, p = 0.103 | 0.471 | 0.215 | 0.908 |
| Eotaxin | F (1, 88) = 0.0001217, p = 0.991 | F (1, 88) = 0.1001, p = 0.752 | F (1, 88) = 5.265, p = 0.024 | 0.994 | 0.313 | 0.267 |
| CCL3 | F (1, 83) = 0.3499, p = 0.556 | F (1, 83) = 3.218, p = 0.077 | F (1, 83) = 0.1847, p = 0.669 | 0.277 | 0.867 | 0.999 |
| CCL4 | F (1, 87) = 0.04708, p = 0.829 | F (1, 87) = 0.5210, p = 0.472 | F (1, 87) = 3.278, p = 0.074 | 0.883 | 0.62 | 0.372 |
| CXCL10 | F (1, 84) = 0.03293, p = 0.856 | F (1, 84) = 0.5949, p = 0.443 | F (1, 84) = 2.615, p = 0.110 | 0.873 | 0.683 | 0.492 |
| G-CSF | F (1, 83) = 0.002348, p = 0.962 | F (1, 83) = 0.02233, p = 0.882 | F (1, 83) = 0.2775, p = 0.600 | 1 | 0.983 | 0.967 |
| PDGF-BB | F (1, 76) = 0.7406, p = 0.392 | F (1, 76) = 2.693, p = 0.105 | F (1, 76) = 1.754, p = 0.189 | 0.234 | 0.984 | 0.311 |

**Supplementary Table 2.** P values from a two-way ANOVA with Šídák Multiple Comparison Test for serum complement and cytokine concentrations in drug-resistant epilepsy (DRE) and healthy groups. Categorical variables: condition [DRE or healthy control (H)] and sex [female (F) or male (M)].
